# Transcription start site analysis reveals widespread divergent transcription in *D. melanogaster* and core promoter-encoded enhancer activities

**DOI:** 10.1101/221952

**Authors:** Sarah Rennie, Maria Dalby, Marta Lloret-Llinares, Stylianos Bakoulis, Christian Dalager Vaagensø, Torben Heick Jensen, Robin Andersson

## Abstract

Mammalian gene promoters and enhancers share many properties. They are composed of a unified promoter architecture of divergent transcripton initiation and gene promoters may exhibit enhancer function. However, it is currently unclear how expression strength of a regulatory element relates to its enhancer strength and if the unifying architecture is conserved across Metazoa. Here we investigate the transcription initiation landscape and its associated RNA decay in *D. melanogaster*. Surprisingly, we find that the majority of active gene-distal enhancers and a considerable fraction of gene promoters are divergently transcribed. We observe quantitative relationships between enhancer potential, expression level and core promoter strength, providing an explanation for indirectly related histone modifications that are reflecting expression levels. Lowly abundant unstable RNAs initiated from weak core promoters are key characteristics of gene-distal developmental enhancers, while the housekeeping enhancer strengths of gene promoters reflect their expression strengths. The different layers of regulation mediated by gene-distal enhancers and gene promoters are also reflected in chromatin interaction data. Our results suggest a unified promoter architecture of many *D. melanogaster* regulatory elements, that is universal across Metazoa, whose regulatory functions seem to be related to their core promoter elements.

## Introduction

Spatio-temporal control of metazoan gene expression is mediated in part by factors acting at gene promoters and at gene-distal transcriptional enhancers. Although major efforts have been made to identify the locations of transcriptional regulatory elements (TREs, here denoting enhancers and promoters) and their cell type-restricted activities, the regulatory mechanisms of these genomic regions are not well understood. Careful characterisation of the properties of TREs and the determinants of their regulatory activity is crucial to better understand the means by which cells control gene expression. Despite the often adopted view on enhancers and gene promoters as distinct entities with discernible functions and local chromatin characteristics (Creyghton et al., 2010; Heintzman et al., 2009, 2007), e.g. different levels of H3K4me1, H3K4me3 and H3K27ac at nucleosomes flanking TREs, recent observations suggest large similarities between mammalian enhancers and gene promoters (Andersson, 2015; Andersson et al., 2015b; Core et al., 2014; Kim and Shiekhattar, 2015). In particular, mammalian TREs are characterised by a high prevalence of divergent transcription initiation (Andersson et al., 2015a, 2014a, b; Kim et al., 2010; Preker et al., 2008; Seila et al., 2008; Sigova et al., 2013). In addition, enhancers frequently contain core promoter elements (Andersson et al., 2014a; Core et al., 2014), bind general transcription factors (GTFs) (Koch et al., 2011; Liu et al., 2011; Zhou et al., 2013), and may act as alternative gene promoters (Kowalczyk et al., 2012). Gene promoters themselves form stable chromatin interactions with other promoters (Chepelev et al., 2012; Li et al., 2012), often resulting in the co-expression of genes in a tissue-specific manner (Chepelev et al., 2012). Several examples of mammalian gene promoters exhibiting enhancer function have also been identified (Dao et al., 2017; Engreitz et al., 2016; Leung et al., 2015; Li et al., 2012). Taken together, these observations raise the question whether the repertoire of TREs may be treated as a unified class (Andersson, 2015; Andersson et al., 2015b; Core et al., 2014; Kim and Shiekhattar, 2015). However, it is currently unclear how promoter (expression) strength relates to enhancer strength, whether a TRE with strong enhancer function also possesses strong promoter function or vice versa, or if enhancer function is inversely related to promoter function. In addition, the inherent state of divergent transcription at TREs in Mammalia is so far not well supported across Metazoa. Observations in *D. melanogaster* have suggested a distinct reduction in divergent events at gene promoters (Core et al., 2012; Nechaev et al., 2010) and less widespread occurrences of enhancer transcription (Core et al., 2012). These observations raise the question whether divergent transcription and, hence, the unifying promoter architecture across TREs is not a conserved property across Metazoa.

Transcription of mammalian protein-coding genes into mRNA is coupled with upstream transcription in the reverse orientation (Andersson et al., 2015a, 2014b; Preker et al., 2008; Seila et al., 2008; Sigova et al., 2013). The latter results in relatively short, non-coding transcripts, commonly referred to as promoter upstream transcripts (PROMPTs) or upstream antisense RNAs (uaRNAs) (Preker et al., 2008; Seila et al., 2008). While core promoters in general possess unidirectional transcription initiation (Duttke et al., 2015), divergent transcription is accomplished by a pair of divergent core promoters contained within the same nucleosome deficient region (NDR) (Andersson et al., 2015a; Scruggs et al., 2015). Divergent transcription initiation is also widespread at regulatory active mammalian transcriptional enhancers (Andersson et al., 2014a; Kim et al., 2010). The resulting transcripts, known as enhancer RNAs (eRNAs), are, similar to PROMPTs, short, low abundant and non-coding. Both PROMPTs and eRNAs are unstable and targets of the ribonucleolytic RNA exosome complex (Andersen et al., 2013; Andersson et al., 2014a, b; Ntini et al., 2013; Preker et al., 2008). The path to RNA decay is, at least in part, linked to the presence of early polyadenylation sites and a depletion of U1 small nuclear ribonucleoprotein (snRNP) binding via a lack of 5’ splice sites downstream of PROMPT and eRNA transcription start sites (TSSs), leading to early transcriptional termination (Almada et al., 2013; Ntini et al., 2013). Like PROMPTS, eRNAs are of low abundance and seldom transcribed from evolutionary constrained DNA (Andersson et al., 2014b; Marques et al., 2013), indicating high similarities between RNA species and that both PROMPTs and eRNAs likely possess little functional relevance. Nevertheless, promoter activity of mammalian enhancers, as observed from local transcription initiation events, is an accurate indicator of active enhancer regulatory function (Andersson et al., 2014a), suggesting a link between distal regulatory enhancer function and local promoter activity. A notable difference between gene promoters and gene-distal enhancers is that while the transcriptional activity at mRNA promoters is strongly favouring the production of stable, exosome-insensitive RNAs on the sense strand, enhancers are generally associated with more balanced production of unstable eRNAs on both strands (Andersson et al., 2014a, b; Core et al., 2014). The apparent commonalities and differences in transcription initiation patterns, frequencies, and RNA decay have therefore been utilised for classification of TRE function (Andersson et al., 2014a, b; Core et al., 2014).

Efforts to catalogue genome-wide the enhancer potential of genomic sequences for activating transcription at a given core promoter have provided insights into the regulatory potential of *D. melanogaster* genomic sequences. Assays based on self-transcribing regulatory regions (STARR-seq) (Arnold et al., 2013), have revealed apparent differences between housekeeping and developmental enhancer activities, as measured by STARR-seq constructs containing core promoters associated with broad, housekeeping activities (hkCPs) and cell type-restricted or developmental core promoters (dCPs), respectively. Sequences activating the former appear to be gene promoter-proximal while sequences activating the latter are generally gene promoter-distal (Zabidi et al., 2015). However, the DNA sequence itself is not the only determinant of enhancer activity. A recent study utilising massively parallel reporter assays emphasizes the importance of the positioning of TREs within chromatin contexts (Corrales et al., 2017). In addition, the boundaries of *D. melanogaster* topologically associating domains (TADs) (Cubeñas-Potts et al., 2017; Sexton et al., 2012) may constrain sequence-compatible enhancer-promoter regulation. Concomitantly, enhancer classes, as characterised by STARR-seq, follow distinct chromatin architectures with respect to TADs, with housekeeping enhancers enriched at domain borders and developmental enhancers enriched at the anchors of loops (Cubeñas-Potts et al., 2017). These results are supported by a preferential enrichment of housekeeping gene promoters at TAD boundaries (Hug et al., 2017).

In this study, we set out to investigate the link between promoter activity and enhancer function and how invertebrate and mammalian genomes compare in their RNA decay and transcription initiation frequencies at TREs. To this end we measured TSS usage, RNA abundance and exosome sensitivity in *D. melanogaster* to assess whether properties (abundance, stability, directionality, divergent transcription) of TREs are conserved across Metazoa and what determines their transcriptional activities. Surprisingly, we find that divergent transcription is a common state of *D. melanogaster* gene promoters and gene-distal enhancers. Characterisation of open chromatin loci into major classes, unbiased to gene annotation, solely by their transcriptional properties recapitulates mammalian archetypical groupings, which reflect gene annotations and enhancer potentials. We show that fly TREs carry remarkable similarities in terms of promoter functionality, regardless of type, pointing to a unified architecture of TREs that is similar across Metazoa. We identify quantitative relationships between TRE expression level and enhancer function, which seem to be encoded by core promoter element strengths, pointing at a regulatory trade-off between developmental enhancer function and promoter functionality and a joint encoding of promoter and enhancer functionality for housekeeping TREs. Our results further suggest at least two layers of transcription control, which are also supported by chromatin interaction data. One, in which housekeeping gene promoters act as enhancers to other gene promoters alike and one in which gene-distal developmental enhancers control the transcription of developmental genes.

## Results

### Fly regulatory elements are associated with divergent transcription and RNA species-specific decay

To characterise transcription initiation events in *D. melanogaster,* we performed deep Cap Analysis of Gene Expression (CAGE (Takahashi et al., 2012)) sequencing (33.5-46.4 million mapped reads per library) in Schneider line 2 (S2) cells. To measure exosome sensitivity, cells were subjected to a double knockdown of the catalytic subunits Dis3 and Rrp6 of the ribonucleolytic exosome complex (by RNA interference, Methods). This resulted in a marked reduction in the abundances of tags aggregating at the annotated TSSs of *Dis3* and *Rrp6* genes when compared to control (dsRNA against GFP) libraries (Supplementary Fig. S1A). The great majority of CAGE tags were proximal to open chromatin regions as measured by DNase I hypersensitivity (82%-86% within 500 base pairs (bp), Supplementary Fig. S1B), indicating a high signal-to-noise ratio of the mapped CAGE data. We observed several instances of divergent transcription initiation, with exosome-sensitive PROMPTs originating upstream of FlyBase (Gramates et al., 2017) gene TSSs in a divergent manner and in a good agreement between replicates (exemplified by Fig. 1A). In addition, many enhancers were associated with replicate-consistent exosome-sensitive divergent eRNAs (exemplified by Fig. 1B). These observations suggest that fly biogenesis and decay of eRNAs and PROMPTs may match those of human.

To quantify the extent and characteristics of divergent transcription in fly cells, we clustered proximally mapped CAGE tags into genomic regions representing CAGE-inferred TSSs (referred to as tag clusters (TCs)). Wide TCs were trimmed and those representing multi-modal peaks were split into narrow single-peak TCs (Supplementary Methods). Although TC expression levels were largely concordant between biological replicates (Supplementary Figs. S2, S3), we filtered out low-level expression (not statistically indistinguishable from genomic background noise, see Supplementary Methods), allowing us to accurately assess the transcriptional patterns of TREs naturally associated with low abundant RNAs (like eRNAs and PROMPTs). This resulted in a total of 147,379 TCs significantly expressed in at least two exosome knockdown replicates. For each filtered TC, we measured the fraction of expression in knockdown conditions to that observed in control libraries, providing a quantitative measure of exosome sensitivity ranging between expression levels fully captured by control CAGE data (exosome sensitivity 0) to expression levels only observed upon exosome knockdown (exosome sensitivity 1). Overall, the majority of TCs associated with annotated mRNA TSSs (~62%) displayed low (<0.25) exosome sensitivity (Fig. 1C). In contrast, a large fraction of PROMPTs (TCs <500 bp upstream of and antisense to annotated FlyBase gene TSSs), lncRNAs (TCs associated with annotated FlyBase ncRNA TSSs) and eRNAs (TCs associated with gene TSS-distal dCP STARR-seq enhancers) were mainly highly (>0.75) exosome sensitive (~51%, ~42%, and ~60%, respectively).

We next calculated the distance from each plus strand TC to the nearest upstream (non-overlapping) minus strand TC (Fig. 1D). ~47% and ~60% of CAGE TCs had, regardless of annotation, a nearest upstream minus strand TC within 250bp and 500bp, respectively. The relative increase in the divergent fraction was reduced at larger distances, suggesting that many fly divergent events are contained within the same NDR (that are most often <500 bp in size). Among specific TREs, we observed that transcribed gene-distal dCP enhancers were frequently (~81%) divergently transcribed, which is supported by a recent study based on nascent RNA sequencing (Meers et al., 2017). Proximal bidirectional (head-to-head) gene pairs showed the highest degree of divergent transcription (~90%), often explained by paired gene TSS usage (data not shown). Stand-alone mRNA promoters, on the other hand, exhibited the least degree of divergent transcription (~46%). These fractions are likely underestimates, since rare transcripts can fall below imposed noise thresholds. Nevertheless, these results demonstrate that a considerable proportion of transcription initiation events in *D. melanogaster* are divergent, and that a fraction of these events are associated with exosomal RNA decay, in accordance with human cells.

**Figure 1.**
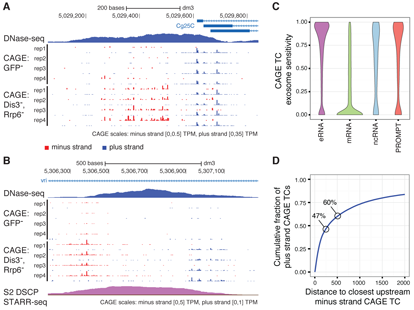
Fly regulatory elements are associated with divergent transcription initiation. **A,B:** Genome browser views around FlyBase annotated TSSs of the *Cg25C* (also known as *Col4a1*) gene (A) and a *vri* intragenic dCP STARR-seq enhancer (B). DNase-seq, control CAGE, and exosome KD CAGE data are shown. All four replicates per CAGE condition are displayed (red: minus strand, blue: plus strand). For visibility reasons, the scales of CAGE signal differ between strands and are provided below each panel. See Supplementary Figures S6 and S7 for RT-qPCR validations. **C**: Distributions of exosome sensitivity, ranging between 0 (insensitive) to 1 (CAGE expression not observed without exosome KD), for eRNAs (associated with dCP STARR-seq enhancers), FlyBase mRNAs, FlyBase ncRNAs and PROMPTs (upstream of and antisense to annotated FlyBase gene TSSs). **D**: Cumulative fraction (vertical axis) of plus strand CAGE TCs that are within a certain distance (horisontal axis) of minus strand CAGE TCs. The percentages of divergent events are highlighted for distances of 250 and 500 bp.

### Expression and exosome sensitivity patterns characterise distinct regulatory elements

With an aim to systematically characterise transcription initiation events and associated RNA turnover at TREs unbiased to existing annotations, we focussed the remaining analyses on DNase I hypersensitive sites (DHSs) (Methods). We quantified DHS-associated expression through aggregation of CAGE tags in strand-specific windows of 200 bp immediately flanking DHS centre points that maximised CAGE tag coverage (Supplementary Fig. S4, Methods). 9,471 out of 11,947 (~79%) called DHSs were significantly expressed (above estimated background noise levels) on any strand in at least two control or exosome depleted libraries. Below, we refer to the most highly expressed strand from a DHS as the ‘major’ strand and the other strand as the ‘minor’. For each DHS, we quantified expression-associated properties on a per-replicate basis according to the knockdown-ascertained major and minor strand expression levels, major and minor strand exosome sensitivity scores, and exosome knockdown-derived transcriptional directionality (Methods). We then performed unsupervised clustering of the transcriptional properties from each of the four replicates across transcribed DHSs, based on a two step clustering procedure (Methods). First we clustered all DHSs into six groups, regardless of replicate. We then compared the resulting group allocation across the replicates for each DHS and clustered a second time, thus generating a final set of clusters, which strongly agreed per DHS across replicates (Supplementary Fig. S5). This resulted in six major groups (Fig. 2A), each of which disagreed on average by at most one replicate within-group (Supplementary Table S1). Only 15 DHSs were removed from further analyses due to lack of replicate agreement, demonstrating that inferred DHS groupings are robust against biological replicate variance. Results from CAGE data were further validated across randomly selected loci by RT-qPCR (Supplementary Figs. S6, S7, Supplementary Table S2), demonstrating the accuracy in determining RNA abundance and turnover even at lowly expressed TSSs.

The clustering of DHSs revealed several interesting relationships between expression levels, transcriptional directionality and exosome sensitivity, and displayed widely different enrichments of annotated gene TSS proximities (Fig. 2B, Supplementary File 1). Three identified classes of DHSs were associated with stable (exosome insensitive) RNAs on their major strand. The proximal regions of *unidirectional stable* and *unidirectional stable w*/ *PROMPT* DHSs were highly enriched (*P* < 2.2 × 10^−16^, Chi-squared test) with annotated unidirectional gene TSSs, consistent with a strong directional expression bias resulting from high expression levels and low exosome senstivity from their major strands. In contrast to *unidirectional stable* DHSs, DHSs in the *unidirectional stable w*/ *PROMPT* category were in addition associated with lowly expressed and highly unstable RNAs from their minor strands, properties reminiscent of human mRNA gene promoters associated with PROMPT transcription (Andersen et al., 2013; Andersson et al., 2014a, b; Ntini et al., 2013; Preker et al., 2008). We also identified a smaller class with more balanced, stable, high expression on both strands (*bidirectional stable*). DHSs in this class were enriched (*P* < 2.2 × 10^−16^, Fisher’s exact test) in annotated head-to-head gene TSSs. We collectively refer to these three classes as *stable* TREs, due to the low exosome sensitivity of RNAs transcribed from their major strands.

The remaining DHSs were grouped into three classes associated with exosome-sensitive RNAs emitted from their major strands (*unstable* TREs, Fig. 2A). One class (*weak bidirectional unstable*) gathered DHSs associated with balanced low output of unstable RNAs. *Intermediate bidirectional stable* DHSs exhibited moderately higher expression on the major strand resulting in a more biased directional transcription. DHSs having close to unidirectional output of unstable RNAs were grouped in the third class of *unstable* DHSs (*weak unidirectional unstable*). All three unstable clusters were highly enriched (*P* < 2.2 × 10^−16^, Chi-squared test) in gene TSS-distal regions (Fig. 2B).

We next compared DHS classes by their association with genome-wide signals from STARR-seq data using constructs based on housekeeping (hkCP) and developmental (dCP) core promoter types (*RpS12* and *even skipped* core promoters, respectively) (Zabidi et al., 2015). Thresholding the log_2_ fold change between STARR-seq signal and input (>1.5) allowed us to compare the number of overlaps with transcribed DHSs (Fig. 2C). Overall, as many as ~65% of transcribed DHSs overlapped a STARR-seq positive region. Specifically, with the exception of *weak unidirectional unstable* DHSs, *unstable* DHSs were enriched (*P* < 2.2 × 10^−16^, Chi-squared test) in dCP enhancers, with the largest overlap (~55%) found among *weak bidirectional unstable* DHSs. Importantly, these results provide external evidence that balanced bidirectional output of exosome-sensitive RNAs is a marker of gene promoter-distal enhancers in *D. melanogaster,* which has previously been established in human cells (Andersson et al., 2014a, b; Core et al., 2014). Interestingly, in agreement with previous reports (Zabidi et al., 2015), a large fraction of *stable* DHSs overlapped with hkCP positive enhancers. The largest overlaps were observed for *bidirectional stable* (~80%) and *unidirectional stable* (~71%) DHSs. *Weak unidirectional unstable* DHSs had modest overlap with STARR-seq positive enhancers and displayed no real preference to either housekeeping or developmental core promoters.

**Figure 2.**
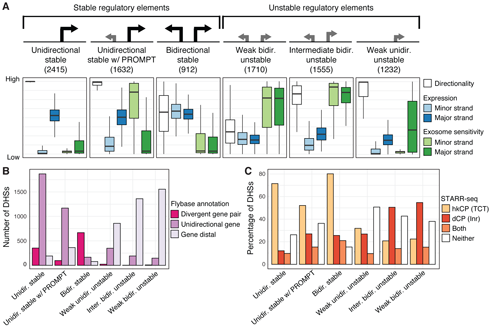
Transcriptional directionality, expression level and exosome sensitivity reveal major groupings of *D. melanogaster* regulatory elements. **A:** Manual labelling and properties of the six identified clusters of transcribed DHSs with similar transcriptional directionality, expression levels, and exosome sensitivities (displayed in box-and-whisker plots). DHS clusters associated with stable or unstable RNAs on their major strands are indicated above DHS cluster labels. See Supplementary Figure S4 for a schematic of the measures used for clustering and the strategy behind expression quantification of DHSs. The lower and upper hinges of boxes correspond to the first and third quartiles of data, respectively, and the whiskers extends to the largest and smallest data points no further away than 1.5 times the interquartile range. For improved visibility, outlier data points are not visualised. **B**: The number of DHSs in each cluster that are in close proximity with divergent (head-to-head) FlyBase gene TSS pairs (divergent gene pair), stand-alone FlyBase gene TSSs (unidirectional gene), or distal from FlyBase gene TSSs. **C**: The percentage of DHSs in each cluster that are overlapping with or are distal to called STARR-seq enhancers, broken up by those overlapping with hkCP enhancers, dCP enhancers or both classes.

In conclusion, clustering of DHSs by their transcription initiation frequencies and associated exosome sensitivity reveals overall similarities between derived clusters of DHSs in *D. melanogaster* S2 cells and those identified in human cells (Andersson et al., 2014b). In addition, a large proportion of *D. melanogaster* TREs show archetypal mammalian-derived properties of PROMPTs and eRNAs. This suggests that mammalian and invertebrate genomes share similar classes and promoter architectures of TREs.

### DNA sequence elements reflect transcriptional directionality and RNA instability

The differences in annotation preferences and transcriptional directionalities between *stable* and *unstable* DHSs prompted us to investigate the relationships between transcriptional output (directionality and RNA exosome sensitivity) and DNA sequence elements at the core promoters and in regions downstream of TSSs of transcribed DHSs. First, we assessed the frequencies of predicted 5’ splice sites and termination signals (polyadenylation sites) at the locations of minor and major strand CAGE summits and up to 1,000 bp downstream (Fig. 3A,B, Methods). Similar to what has been previously observed in human (Almada et al., 2013; Ntini et al., 2013), we observed an enrichment in downstream 5’ splice sites on the major strands of *stable* DHSs while site frequencies were close to or under the genomic background level for their minor strands and for both strands of *unstable* DHSs (Fig. 3A), indicating that the instability of RNA is inversely related to downstream flanking 5’ splice site sequences. Enrichments of 5’ splice sites were supported by a higher prevalence of multi-exonic transcripts, inferred from RNA-seq data (Lim et al.), arising from the major strands of *stable* DHSs (Fig. 3C). In contrast, unstable RNAs were mostly unspliced. In further agreement with the human system, we noted that polyadenylation sites (AWTAAA consensus hexamers) were in general depleted downstream of stable RNA TSSs, but above or at genomic background levels for unstable RNAs (Fig. 3B). However, in contrast to human (Andersson et al., 2014b), we found an enrichment of polyadenylation sites in the immediate region (within 100 bp) downstream of stable RNA TSSs before falling below the genomic background further (>200 bp) downstream. High frequencies of polyadenylation sites in the 5’ untranslated region (UTR) of many fly mRNAs have previously been characterised (Guo et al., 2011). These enrichment differences between fly and human might reflect differences in sequence preferences between fly and mammalian gene promoters (Kadonaga, 2011; Lenhard et al., 2012), such as a depletion of CpG islands in the *D. melanogaster* genome.

Next, we investigated the prevalence of core promoter elements on minor and major strands of transcribed DHSs (Supplementary File 2). We focused on eight functional core promoter elements (Kadonaga, 2011; Lenhard et al., 2012; Ohler et al., 2002; Rach et al., 2009) and the Trl element (GAGA motif of *Trithorax*-*like*) that, based on motif finding, either had clear preferences for expected positions (TATA, Inr, DPE, MTE (Ohler10), and E-box (Ohler5), Supplementary Fig. S8) or an enrichment in individual DHS classes compared to random genomic background regions distal to DHSs but with weaker positional bias (Ohler1, DRE, Ohler6, and Trl, one-sided Mann-Whitney U test *P* < 1 × 10^−20^). Overall, TATA, Inr, DPE, and MTE elements displayed the strongest positional preferences on major strands among investigated motifs (Fig. 3D). Among the nine motifs, DRE, Ohler6, TATA and Trl elements showed the highest presence on minor strands (Supplementary Fig. S8), although at lower frequencies than on major strands, suggesting that TREs of *D. melanogaster* may be composed of two divergent core promoters. Indeed, a considerable fraction of DHSs were associated with at least one (any) significant core promoter element motif match on both strands (ranging between ~20% for *weak bidirectional unstable* DHSs to ~52% for *bidirectional stable* DHSs, Fig. 3E). In comparison, ~12% of random genomic background regions had significant motif matches on both strands. While these results reflect a potential of having two divergent core promoters across TREs, calling of motif instances in genomic sequences can be inexact and does not directly reflect the strengths of considered core promoter elements.

**Figure 3.**
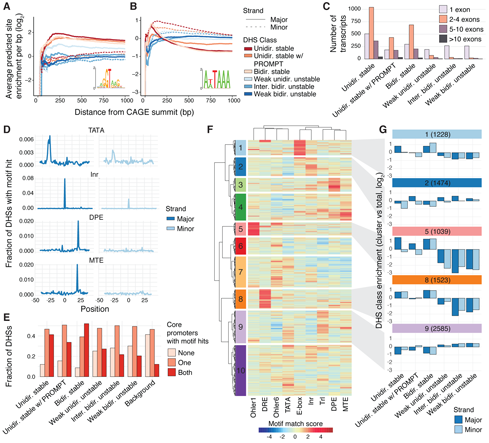
DNA sequence elements impact the stability, directionality and expression strengths of regulatory elements. **A,B**: Frequencies of RNA processing motifs (A: 5’ splice site, B: polyadenylation site) downstream of CAGE summits broken up by DHS class and strand. Vertical axes show the average number of predicted sites per kb within an increasing window size from the TSS (horizontal axis) in which the motif search was done. 0 indicates the expected hit frequency from random genomic background. **C:** Histograms of de novo-assembled transcript counts (vertical axis), broken up by number of exons, associated DHS class and strand. **D:** Fraction of transcribed DHSs (vertical axis) with an identified core promoter element (TATA, Inr, DPE, or MTE) at a given position relative to the major (left panels) and minor (right panels) strand CAGE summits. **E:** Fraction of transcribed DHSs within each DHS class (vertical axis) associated with at least one out of nine core promoter elements identified on one or both strands. In adition the fraction of DHSs with no core promoter elements are shown (None). **F:** Hierarchical Ward agglomerative clustering of motif match scores for the nine considered core promoter elements on major and minor strands of transcribed DHSs. Ten clusters of core promoter element compositions are shown. **G:** DHS class enrichments, calculated as the fraction of DHSs in each DHS class associated with each core promoter element cluster versus the fraction of total transcribed DHSs, displayed in log_2_ scale enrichment, broken up by major and minor strand. See Supplementary Figure S9 for DHS class enrichments for all core promoter clusters.

To investigate potential differences between DHS classes as well as between minor and major strands, taking into account the strengths of motif matches, we clustered the maximum match scores for each considered core promoter element motif in regions surrounding the CAGE summits of minor and major strands (Methods). This revealed a complex combination of core promoter elements across DHSs, for which many core promoter elements were restricted to only a subset of DHS regions (Fig. 3F), reflecting the wide diversity of core promoter compositions in *D. melanogaster* (Kadonaga, 2011; Ohler et al., 2002). Individual core promoter clusters displayed strong match scores for individual core promoter elements, such as E-box (cluster 1), Inr (cluster 2), DPE (cluster 3), MTE (cluster 4), Ohler1 (cluster 5), DRE (cluster 8), and Trl (cluster 9). We observed several preferences between core promoter elements and the stability of associated RNAs (Fig. 3G, Supplementary Fig. S9A). For instance, cluster 5 (associated mainly with Ohler1) had a strong enrichment of major strands of *unidirectional stable* and *unidirectional stable w*/*PROMPTs* DHSs (Fisher’s exact test, *P* < 1 × 10^−16^, *P* < 1 × 10^−10^, respectively), as well as both strands of *bidirectional stable* DHSs (Fisher’s exact test, *P* < 1 × 10^−16^, both), but a strong depletion of core promoters of *unstable* DHSs (Fisher’s exact test, *P* < 1 × 10^−10^). In contrast, Trl elements (cluster 9) were mostly found in core promoters of *unstable* DHSs. Other core promoter clusters were not mainly associated with the stability of produced RNAs. In particular, Inr elements (mostly identified in cluster 2) showed no clear differences between *stable* and *unstable* DHSs, but rather a preference for major over minor strands. Interestingly, cluster 8 (associated mainly with DRE) was enriched with *unidirectional stable* and *bidirectional stable* DHSs (Fisher’s exact test, *P* < 1 × 10^−15^, all) but not *unidirectional stable w/PROMPTs* DHSs. In general, we found that *bidirectional stable* and *unidirectional stable* major strands showed similar enrichment within core promoter clusters, while *unidirectional stable w*/*PROMPTs* major and minor strands showed higher similarities with *unstable* DHS groups (Supplementary Fig. S9B).

Taken together, we conclude that *D. melanogaster* TREs are frequently associated with core promoter elements regardless of DHS class but possess strong diversity in core promoter composition that are reflecting RNA stability and transcriptional directionality.

### Genomic positioning and core promoter elements may impede divergent transcription

We next wanted to investigate the nature of absent PROMPT transcription from *unidirectional stable* DHSs. Invertebrate genomes have an unexpectedly high fraction of head-to-head gene pairs, not immediately explained by their more compact genomes compared to mammalian ones (Yang and Yu, 2009). Given that the distance between head-to-head gene pair TSSs has an observable impact on human PROMPT transcription (Chen et al., 2016), we compared the distances between upstream antisense gene TSSs and major strand CAGE summits of *unidirectional stable* and *unidirectional stable w*/ *PROMPTs* DHSs. We noted a clear difference in positional preference between DHS classes (Fig. 4A). Major strand TSSs of *unidirectional stable* DHS were more frequently positioned in close proximity (within 1,000 bp) of upstream annotated head-to-head gene TSSs than the major strand TSSs of *unidirectional stable w*/ *PROMPTs* DHSs (Fisher’s exact test, *P* < 2.2 × 10 ^−16^). In support, we found that plus strand TCs (regardless of DHS class) positioned within 500-1,000 bp from minus strand gene TSSs were more frequently associated with unidirectional than divergent transcription (evaluated by the frequency of divergent events within 500 bp), while the divergent fraction increased at distances >1,000 bp (Fig. 4B). In contrast, at distances below 500 bp most transcription events were divergent. Hence, PROMPT transcription seems to be impeded when the promoter is placed in close proximity (within 1,000 bp) with other gene TSSs in a head-to-head orientation. However, a considerable fraction of *unidirectional stable* DHS could not be explained solely by distance constraints (~40% of such DHSs are >2,000 bp from upstream head-to-head gene TSSs).

**Figure 4.**
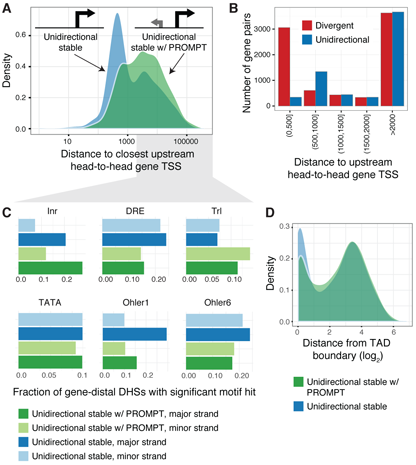
PROMPT transcription is impeded by positional and core promoter element constraints. **A**: Densities of the distances between DHS major strand CAGE summits and the closest upstream antisense FlyBase gene TSS (head-to-head composition) for *unidirectional stable* and *unidirectional stable w*/ *PROMPTs* DHSs. **B**: The number of divergent and unidirectional events (vertical axis) for CAGE TCs at various distances from the closest upstream antisense FlyBase gene TSS (head-to-head composition). Divergent events were defined as divergent TC summits within 500 bp. **C**: Fraction of *unidirectional stable* and *unidirectional stable w*/ *PROMPTs* DHSs positioned >2,000 bp from the closest upstream antisense FlyBase gene TSS having core promoter elements Inr, DRE, Trl, TATA, Ohler1, and Ohler6 on major and minor strands. **D**: Densities (vertical axis) of distances between transcribed DHSs to TAD boundaries (horisontal axis) for *unidirectional stable* and *unidirectional stable w*/ *PROMPT* DHSs.

At distances >2,000 bp to upstream head-to-head gene TSSs, we observed notable sequence differences between *unidirectional stable* DHSs compared to those with PROMPTs (Fig. 4C). In particular, Ohler1 and DRE core promoter elements on the major strand were more frequently associated with unidirectional DHSs than those with PROMPTs (Fisher’s exact test, Ohler1: *P* < 3.5 × 10^−16^, DRE: *P* < 4.3 × 10^−8^). Overall, core promoter clusters (Fig. 3F) defined by these elements (clusters 5 (Ohler1) and 8 (DRE)) were associated with a higher transcriptional directionality score (Supplementary Fig. S9C). In contrast, Trl element occurrence was associated with PROMPT transcription (Fig. 4C) and the core promoter cluster (cluster 9) most strongly associated with this element displayed the weakest transcriptional directionality scores (Supplementary Fig. S9C). Interestingly, both Ohler1 and DRE elements are associated with broader, ubiquitous expression (Lenhard et al., 2012), while Trl elements do associate with regulated, cell type-constrained gene expression (Zabidi et al., 2015). In addition, it is known that DRE and Trl elements have different positional preferences in chromatin architectures, with DRE elements frequently co-occurring with housekeeping TREs at TAD boundaries (Cubeñas-Potts et al., 2017). In agreement, *unidirectional stable* DHSs were more frequently positioned in the vicinity of TAD boundaries (Cubeñas-Potts et al., 2017) than *unidirectional stable w*/ *PROMPTs* DHSs (Fig. 4D) (Fisher’s exact test, *P* = 1.297 × 10^−9^).

In summary, the prevalence of divergent transcription in *D. melanogaster* may be impeded by constraints on core promoter element composition, genomic positioning, and proximal chromatin architectures. Since many of these features differ in frequencies from mammalian genomes, we suggest that these characteristics explain the lower tendency of divergent transcription at *D. melanogaster* gene promoters.

### Enhancer potential is related to endogenous expression level

Next, we investigated the association between chromatin state and DHS class. We overlaid transcribed DHSs with locations of histone modifications, histone variants, and TF binding sites (binarised modENCODE (modENCODE Consortium et al., 2010) ChIP-chip data (Zhou and Troyanskaya, 2016)), and calculated the per-class binding proportions within 100 bp of the major strand CAGE summit (Figure 5A). Several chromatin marks clearly distinguished *unstable* from *stable* DHSs, including H3K4me1, H3K18ac, H4K8ac, H4K5ac, and H3K27ac at *unstable* DHSs, and H2AV and H3K4me3 at *stable* DHSs. Other histone modifications did not specifically follow the inferred stability classes. H3K9ac were mostly found at *bidirectional stable* DHSs, while H2BK5ac displayed a promiscuous association with transcribed DHSs. We observed preferential GAF binding to *unstable* DHSs and to some extent also *unidirectional stable w*/ *PROMPT* DHSs, confirming the observed Trl element enrichment in these DHS classes (Figure 3F,G). Interestingly, architectural proteins frequently residing at chromatin domain boundaries (Sexton et al., 2012; Wang et al., 2017), such as CTCF, BEAF32, CP190, Chriz (Chromator) and its associated kinase JIL1, frequently overlapped *stable* DHSs.

Although H3K4me1 and H3K4me3 showed preferential overlaps with *unstable* and *stable* DHSs in agreement with preferential gene TSS-distal enhancer associations (Fig. 2C), we wanted to investigate their association with STARR-seq enhancers regardless of DHS class. First, we split *bidirectional weak unstable* and *unidirectional stable w*/ *PROMPT* DHSs according to whether they overlapped with a hkCP enhancer or a dCP enhancer and plotted enrichments of H3K4me1 and H3K4me3 along with two other chromatin marks distinguishing *stable* from *unstable* DHSs, namely H3K18ac and H2AZ (Figure 5B, Supplementary Fig. S10). For all four marks, we observed a binding profile which appeared remarkably similar according to the overlapped enhancer class, with binding reflecting the hkCP or dCP enhancer potential as opposed to the DHS class itself. H3K4me1 and H3K18ac were both depleted at the centre of DHSs for both classes when overlapping with hkCP enhancers, whilst enriched for both classes when overlapping dCP enhancers. In contrast, H2AV and H3K4me3 overlaps were frequent in cases for which either DHS class overlapped hkCP enhancers and showed a much reduced frequency around DHSs overlapping dCP enhancers. Interestingly, based on ChIP-seq data (Herz et al., 2012), we observed quantitative relationships (Spearman rank correlation test, *P <* 2.2 × 10^−16^) between H3K4me3 and H3K4me1 levels and enhancer strengths (Fig. 5C,D). In line with preferential enrichments in DHS classes and enhancer classes, H3K4me3 displayed a positive correlation with hkCP enhancer potential (Spearman’s rho = 0.49) and a weaker negative correlation with dCP enhancer strength (Spearman’s rho = −0.26), while H3K4me1 showed the opposite trends. Enhancer strength associations were also strong for the ratio between H3K4me1 and H3K4me3, but interestingly not for H3K27ac (Supplementary Fig. S11). Thus, it appears that H3K4me1 and H3K4me3 levels, as well as the ratio between these, reflect the underlying DNA sequence and in particular the ability of the sequence to act as an enhancer.

However, H3K4me3 and H3K4me1 are both associated with transcriptional levels (Core et al., 2014), and likely reflect transcriptional memory and consistency between cells (Howe et al., 2017). Congruently, H3K4me1:H3K4me3 ratio and H3K4me1 levels (Supplementary Files 3 and 4) were negatively associated with endogenous CAGE expression levels (Fig. 6A and Supplementary Fig. S11, Spearman rank correlation test, *P* < 2.2 × 10^−16^, rho = −0.46). In line with this observation, dCP enhancers were less expressed than DHSs not associated with dCP enhancer potential (t-test, *P* < 1 × 10^−16^), while hkCP enhancer-associated transcribed DHSs were associated with the highest expression levels (t-test, *P* < 1 × 10^−16^). In general, we observed a striking positive correlation between hkCP signal and the major strand CAGE expression level of DHSs regardless of attributed class, indicating that the stronger the housekeeping enhancer potential, the more transcription is observed from the DHS (Fig. 6B, Spearman rank correlation test, *P* < 2.2 × 10^−16^, rho = 0.43). These results argue that the observed chromatin mark enrichment over DHS clusters and enhancer classes might reflect local core promoter strength. Indeed, the strength of DRE elements (as determined by motif match score) was positively correlated with hkCP enhancer potential (Supplementary Fig. S12), which has been reported earlier (Zabidi et al., 2015). Hence, the enhancer potential to activate hkCP core promoters is related to core promoter strength (as observed for DRE), which is itself biased towards *stable* DHSs (Fig. 3F,G). In contrast, endogenous expression levels were lowest for DHSs with the strongest dCP enhancer potential (Fig. 6C, Spearman rank correlation test, *P* < 2.2 × 10^−16^, rho = −0.20), suggesting that dCP enhancer function is incompatible with strong promoter function. Importantly, the overall trends observed for H3K4me1, H3K4me3, and CAGE expression levels versus housekeeping enhancer potential were consistent both for DHSs that were proximal and those that were distal to FlyBase gene TSSs (Supplementary Fig. S13). In addition, gene TSS-proximal DHSs with strong dCP enhancer potential tended to be more lowly expressed than those with weak dCP enhancer potential.

**Figure 5.**
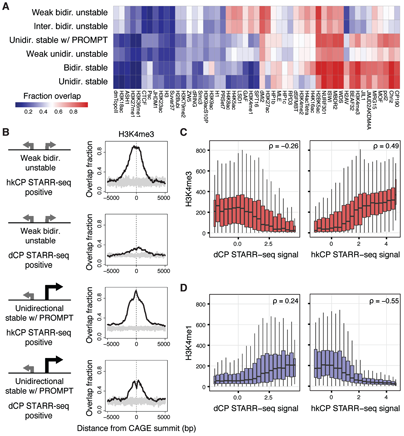
Histone modifications and architectural protein binding reflect enhancer potential. **A**:Heat map representing DHS class proportions of binarised ChIP-chip data. Rows and columns are hierarchically clustered and binding is defined as at least one binding event observed within +/−100 bp of the major strand CAGE summit. **B**: Detailed binding enrichments for H3K4me3 at *weak bidirectional unstable* and *unidirectional stable w*/ *PROMPT* DHSs, broken up according to STARR-seq enhancer potential (overlapping either a hkCP or dCP enhancer), based on binding proportions within 5,000 bp from the CAGE summit. Grey represents background distribution based on randomised locations, generated 10 times per plot. See also Supplementary Figure S10 for the profiles of H3K4me1, H3K18ac, H2Az. **C**: Normalised H3K4me3 ChIP-seq data (vertical axis) versus binned dCP (left) and hkCP (right) STARR-seq signal (horisontal axes). Spearman’s rho statistics calculated on non-binned data are displayed in the top right corners of panels. **D**: Like C but for normalised H3K4me1 ChIP-seq data. Box-and-whisker plots (C,D) displayed as in Figure 2A.

In conclusion, our results suggest that TRE housekeeping enhancer strength is reflecting local core promoter strength and thus, endogenous expression levels. The opposite trend for dCP enhancer potential and local expression level further suggest a regulatory trade-off between promoter strength and developmental enhancer strength.

**Figure 6.**
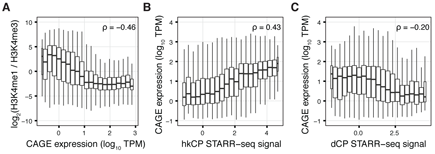
Enhancer potential is related to local endogenous expression levels. **A:**H3K4me1:H3K4me3 normalised ChIP-seq ratio (log_2_, vertical axis) versus DHS major strand CAGE expression levels (log_10_ TPM, horisontal axis). **B:** DHS major strand CAGE expression levels (log_10_ TPM, vertical axis) versus binned hkCP STARR-seq signal (horisontal axis). **C:** Like B, but for binned dCP STARR-seq signal. See also Supplementary Figure S11 for quantitative relationships between expression levels and H3K4me1, H3K4me3, and H3K27ac. Box-and-whisker plots (A-C) displayed as in Figure 2A.

### Three dimensional architectures reveal multiple layers of transcriptional regulation

Next, we utilised TAD information based on high resolution Kc167 HiC data (Cubeñas-Potts et al., 2017) to investigate how defined DHS classes in *D. melanogaster,* with respect to RNA exosome-sensitivity, directionality and expression strength, behave within their three-dimensional contexts. The number of transcribed DHSs within TADs ranged from 1 to 31, with a clear skew towards fewer sites and single-element TADs having the greatest frequencies (Supplementary Fig. S14A). Interestingly, the number of DHSs within a TAD only very weakly correlated with the size of the TAD in which they belonged (Fig. S14B), suggesting that the TREs within multi-element TADs are more densely situated than in TADs with fewer TREs. In general, *stable* DHSs were frequently positioned closer to TAD borders, whereas *unstable* DHSs were often positioned away from boundaries (Supplementary Fig. S15). *Unidirectional stable* DHSs were also more likely to be positioned between annotated TADs as opposed to within (*P* < 2.2 × 10 ^−16^, Fisher’s exact test), while on the contrary, *unstable* DHSs were strongly enriched within the TADs themselves (*P* < 2.2 × 10^−16^, Fisher’s exact test, Fig. 7A).

We asked whether TREs co-localised preferentially with other types of TREs within TADs. Considering TADs containing at least three transcribed DHS, we applied generalised linear models (GLMs) to calculate the odds of a partnering TRE of each DHS class appearing within the same TAD (correcting for the distance between TREs within and between TADs and against a background of encountering a random TRE, see Methods). Overall, the *unidirectional stable* and *bidirectional stable* DHSs showed a preference for co-localisation with DHSs of their own classes, and a reduced preference for *unstable* DHSs (Fig. 7B). Similarly, *unstable* DHSs showed a clear preference for co-localisation with other DHSs of the same category. Interestingly, *unidirectional stable w*/ *PROMPT* DHSs showed a preference towards grouping with the *unstable* DHSs, thus showing a very different trend to its counterpart DHSs without detected PROMPTs.

To investigate if these results reflect a general property or a consequence of multiple layers of regulatory architectures occurring within the genome of *D. melanogaster,* we clustered the TADs according to their memberships of transcribed DHSs (see Methods), generating a total of 7 clusters (Fig. 7C). In agreement with the GLM results, TADs that were highly enriched in *unstable* DHSs also contained *unidirectional stable w*/ *PROMPT* DHSs but very few DHSs classified as *unidirectional stable* or *bidirectional stable* (e.g., clusters 1, 3, 4). In addition, some TADs predominantly contained *stable* TREs (e.g. clusters 6 and 7). The clustering of TAD memberships also revealed other combinations of DHS classes, not apparent from the GLM analysis, e.g. cluster 5, which had a tendency to contain a mixture of classes, in particular combinations of *unidirectional stable* elements with *unstable* DHSs. This cluster might reflect cases in which enhancer-core promoter preferences do not follow our generalised DHS class attributions. In order to investigate the architectural relationship of DHS classes with respect to enhancer potential, we generated models investigating the context of individual chromatin interactions (Cubeñas-Potts et al., 2017) according to the hkCP or dCP enhancer strengths of target DHSs (utilising sequences with no STARR-seq enhancer potential as background). Notably, DHSs associated with *stable* classes interacted preferentially with DHSs possessing hkCP enhancer potential, but *unstable* classes did not show such a preference (Supplementary Fig. S16A), reflecting the strong association between *stable* DHS and local hkCP enhancer potential. Further supporting the separate architectures of dCP and hkCP enhancers and their links with DHS classes, the interaction targets of DHSs possessing dCP enhancer potential were depleted of *stable* DHSs.

Taken together, our results suggest multiple and distinct regulatory architectures for hkCP and dCP enhancers, that can be categorised into two general regulatory programmes. One in which housekeeping gene promoters may act as enhancers of other gene promoters and another in which gene promoters are regulated by gene-distal developmental enhancers, likely affecting cell-type restricted expression levels.

**Figure 7.**
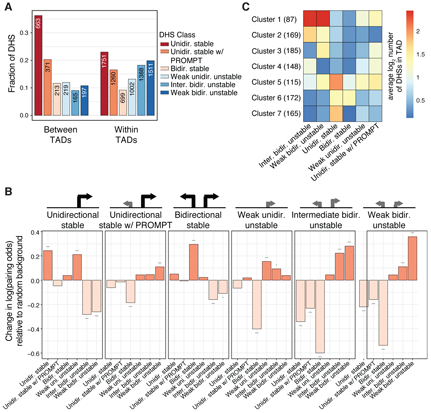
Chromatin architectures suggests multiple layers of transcriptional regulation. **A**:Fractions of DHSs per class, out of either the total within (red) or between (blue) annotated TADs. The number of DHSs in each class is denoted on top of bars. **B**: The change in log(odds) of grouping within the same TAD for DHS classes, split according to grouping class. Significance stars interpreted as: * = *P*< 0.1, ** = *P* < 0.01 or *** = *P* < 0.001. **C**: Heatmap representing clusters of TADs generated according to the membership of TREs defined by DHS classes. Colours represent the average log_2_(number of elements in TAD) for the given DHS class and TAD membership cluster. The number of TADs represented in each TAD cluster is given in parentheses.

## Discussion

In this study we provide an extensive annotation-unbiased characterisation of transcriptional regulatory elements in *D. melanogaster*, based on the biogenesis and properties of produced transcripts, including expression levels, transcriptional directionality and RNA exosome-sensitivity. TSS data of capped RNAs in exosome-depleted S2 cells provide clear evidence that a large fraction of TREs in general are divergently transcribed. In particular, the vast majority of active gene-distal transcriptional enhancers are characterised by divergent transcription of exosome-sensitive eRNAs. In addition, a considerable proportion of gene promoters are associated with divergent transcription of stable mRNAs and unstable PROMPTs. These archetypical properties, distinguishing gene-distal enhancers from gene promoters, allow for accurate classification of regulatory function from expression data alone. We further show that similar principles of downstream RNA processing seen for mammalian PROMPTs and eRNAs are linked to exosome sensitivity also in *D. melanogaster*. Importantly, our observations support a unified divergent promoter architecture for many TREs (Andersson et al., 2015b; Core et al., 2014; Kim and Shiekhattar, 2015), which is similar across Metazoa.

Despite the prevalence of divergent transcription at active TREs, a fraction of gene promoters in *D. melanogaster* are unidirectionally transcribed. We find that genomic positioning and localisation within chromatin architectures might explain some of these exceptions. When a gene TSS is positioned proximal to an upstream gene TSS in a head-to-head conformation, PROMPT transcription is impeded and the distance between divergent events are determined by the distance between paired gene TSSs, lending support to observations in human cells (Chen et al., 2016). Interestingly, divergent gene pair occurrences in invertebrates are much more frequent than what can be explained by their constrained genome size alone (Yang and Yu, 2009). Positioning with respect to TAD structures also seems to have an effect, with unidirectionally transcribed TREs more frequently positioned in close proximity with TAD boundaries than divergent ones. Binding of upstream architectural proteins also seems to have an effect on divergent transcription in human cells (Bornelöv et al., 2015), potentially acting as barriers to elongating RNAPIIs. These features may, at least partially, explain the previously claimed lack of divergent transcription at *D. melanogaster* gene promoters (Core et al., 2012; Meers et al., 2017; Nechaev et al., 2010).

Systematic characterisation of core promoter elements at TREs revealed a complex landscape of core promoter compositions. We observed distinct associations of certain core promoter elements to subsets of TREs, which were strongly associated with their expression strength, directionality and RNA stability. Presence of Trl elements is associated with TREs characterised by balanced bidirectional transcription of unstable RNAs, thereby providing a signature of many gene-distal enhancers. Other elements, including Ohler1 and DRE, are strongly associated with directionality of transcription. Since Ohler1 and DRE are specific to invertebrates (Lenhard et al., 2012), their enrichments provide an additional explanation of the reduced prevalence of PROMPTs at *D. melanogaster* gene promoters compared to mammals.

Integration of enhancer potential data, as measured using STARR-seq assays, provided clear insights into the relationship between transcriptional properties of TREs and the link between promoter and enhancer function. Gene promoter-like loci had a tendency to overlap housekeeping enhancers, as reported previously (Zabidi et al., 2015). Interestingly, we found that housekeeping enhancer strength is strongly related with endogenous CAGE expression levels. This is potentially driven by core promoter element strength, in particular for DRE elements, which itself is correlated with housekeeping enhancer strength. Thus, the stronger the promoter, the greater potential it has to act as a housekeeping enhancer. In contrast, strong developmental enhancers are associated with weak promoter expression levels. Developmental enhancer strength was associated with low endogenous expression levels, suggesting a regulatory trade-off between promoter function and developmental enhancer function. Our identified link between enhancer function, core promoter strength and promoter expression level provides insights into frequently used histone modifications to discern enhancers from gene promoters, including H3K4me1 and H3K4me3, which we find to be related to expression levels. Such histone modifications are therefore likely indirect markers of enhancer function, reflecting the generally weaker promoter strengths and thereby expression levels of gene-distal enhancers. However, although H3K4me1 tended to be prevalent at lowly expressed developmental enhancers, it is important to note that H3K4me1 on its own cannot distinguish between active and inactive enhancers Bonn et al. (2012).

In line with observed differences between developmental and housekeeping enhancers, chromatin conformation data suggest a model involving separate architectures of transcriptional regulation, in which TREs were strongly biased to interact with those with the same transcriptional properties and regulatory potentials. Gene-distal developmental enhancers are enriched within TADs and gene promoters with housekeeping enhancer potential are enriched near TAD borders, reflecting the constraints chromatin architecture can have on transcriptional activity and regulation. Our results suggest at least two distinct regulatory programmes for housekeeping and developmental enhancers. For developmental, or cell type-restricted regulation, gene-distal developmental enhancers seem to regulate gene promoters with cell-type restricted expression levels constrained within the same TAD. Housekeeping gene promoters, on the other hand, are frequently located close to TAD borders, and may act as enhancers to other gene promoters alike.

Importantly, our study implies that enhancers are RNAPII promoters and that regulatory function might not be discernible on a per-element basis. Rather, a fraction of metazoan TREs possesses both strong enhancer and strong promoter function, while others are characterised by strong enhancer function and weak promoter function or vice versa. Based on our observations we favour the most parsimonious model, in which TREs (classically labelled as enhancers or promoters) should be referred to as promoters, whose regulatory activities and effects (local or distal) are determined by local core promoter strength and the genomic landscape and chromatin architecture in which they are placed.

## Materials and Methods

### S2 cell culturing and RNA interference

*D. melanogaster* S2 cells were cultured at 27°C in Schneider’s medium (Sigma, S0146) supplemented with 10% FBS (Sigma, F7524) and 1% penicilin/streptomycin (Sigma, P0781). Double-stranded RNAs (dsRNA) to deplete Rrp6 and Dis3 were prepared by in vitro transcription from a PCR template with T7 promoters on both ends using the Megascript RNAi kit (Ambion, AM1626) according to the manufacturer’s instructions (Supplementary Table S3). DsRNA against GFP was used as a control. For each condition, 3×10^6^ cells were seeded in a well of a 6-well plate. The following day, cells were washed twice with Schneider’s media with no FBS and no antibiotica, a mixture of 40μg of dsRNA (20 μg Rrp6 dsRNA and 20 μg Dis3 dsRNA or 40μg GFP dsRNA) in 500μl media was added dropwise to the cells, the plates were agitated for 30 sec and incubated for 6h at 27°C. Finally, 2.5ml of media with FBS and penicilin/streptomycin were added. The treatment was repeated 2 days later and the cells were harvested 4 days after the first dsRNA treatment.

### CAGE library preparation, sequencing and mapping

CAGE libraries were prepared as described elsewhere (Andersson et al., 2014b; Takahashi et al., 2012) from total RNA purified from S2 cells with TRIzol (Ambion, 15596018) according to the manufacturer’s protocol. Sequenced reads were trimmed to remove linker sequences and subsequently filtered for a minimum sequencing quality of Q30 in 50% of the bases. Mapping to *D. melanogaster* (dm3, r5.33) was performed using Bowtie (Langmead et al., 2009) (version 1.1.1), allowing max two mismatches per read and keeping only uniquely mapped reads.

### Processing of DNase-seq data and identification of DNase I hypersensitive regions

Sequencing reads from DNase-seq and input data (Arnold et al., 2013) were trimmed using Trimmomatic (Bolger et al., 2014) (version 0.32) using a sliding window approach, trimming off the ends of reads in 4 nucleotide windows that did not fulfil a quality ≥Q22. Reads trimmed to a length shorter than 25 nucleotides were discarded. Next, trimmed reads were mapped to the dm3 (r5.33) reference genome using Bowtie (Langmead et al., 2009) (version 1.1.1), allowing max three mismatches per read and keeping only uniquely mapped reads. Next, DNase I hyperensitive sites (DHSs) were called using hotspot (John et al., 2011) at a FDR threshold of 0.01, on input data, pooled DNase-seq data and in each of two DNase-seq replicates. DHS hotspot peaks called from pooled replicates that overlapped peaks from individual replicates and not peaks from input DNase-seq data were used for further analyses. This resulted in a final set of 11,947 DHSs.

### CAGE tag clustering and expression quantification

Tag clustering was performed on pooled CAGE data, including all four replicates from each condition using a strategy to remove tails from wide TCs and split multi-modal peaks (see Supplementary Methods). This resulted in a set of 670,681 TCs.

We next quantified the expression of each TC in each individual CAGE replicate by counting of CAGE 5’ ends falling into their defined genomic regions. In addition, CAGE genomic background noise levels were estimated (see Supplementary Methods). Only TCs whose expression was above the CAGE genomic background noise threshold in at least two out of four replicates in each condition were considered. Noise level filtering resulted in 121,809 TCs in control CAGE libraries and 147,379 TCs in exosome knockdown CAGE libraries. Expression levels were converted to tags per million (TPM), by counting the CAGE tags per TC and normalizing to library size scaled by 10^6^. TCs were annotated to FlyBase gene TSSs based on a max distance between TC summit positions and gene TSS of 250 bp (upstream or downstream).

To measure the effect of the knockdowns, the mean expression and standard deviation over the four replicates for the knockdown and control experiments were calculated for TCs annotated to the primary FlyBase TSSs of the genes *Rrp6* and *Dis3*.

### RT-qPCR validations of CAGE expression levels and exosome sensitivities

Expression levels and exosome sensitivity measured by CAGE were validated using RT-qPCR (Supplementary Table S4). The RNA was treated with TURBO DNA-free kit (Ambion, AM1907) and cDNA was prepared with the SuperScript II kit (Invitrogen, 18064014), using 1μM oligo dT18 and 5ng/μl random primers. qPCRs were performed with Platinum SYBR Green qPCR SuperMix-UDG (Invitrogen, 11744500) in a MX3000P (Agilent technologies) machine. RNA levels were normalised to that of *Act5C.*

### DHSs as focus points for transcription initiation

DHSs were used as focus points for characterising patterns of transcription initiation events at TREs as described elsewhere (Andersson et al., 2014b), with minor modifications (see Supplementary Methods). DHSs were annotated to FlyBase (release 5.12) genes by intersection of gene TSSs with DHSs extended 100 bp upstream and 200 bp downstream with respect to the annotated gene strand. DHSs were subsequently categorised into those associated with gene TSSs on plus strands or minus strands only (unidirectional genes), those associated with both gene TSSs on plus strands and minus strands (divergent gene pairs), or those that were not annotated on any strand (gene TSS-distal).

### Annotation-unbiased clustering of DNase I hypersensitive sites

Unsupervised clustering of DNase I hypersensitive sites was performed on the basis of transcriptional properties derived from each replicate CAGE library in a two-step approach (see Supplementary Methods), to guarantee high agreement between individual replicates.

### Analysis of downstream RNA processing events

To examine the propensity of RNA-processing motifs downstream of TSSs and their association with DHS transcriptional properties, we assessed the occurrences of the 5’ splice site (SS) and polyadenylation site (AWTAAA) termination motifs using HOMER (Heinz et al., 2010). Position weight matrices for 5’ splice sites and polyadenylation sites were collected from the JASPAR database (Mathelier et al., 2016) (ID: SD0001.1, NAME: at_AC_acceptor) and custom-made based on consensus, respectively. The enrichment per bp of motif hits over increasing distances up to 1kb downstream of the CAGE summit, considering the respective strand of each DHS, was calculated by comparison to the predicted motif occurrences from a genomic uniform distribution (enrichment over genomic background).

### Transcript assembly from RNA-sequencing data

Annotation of transcripts was performed by *de novo* assembly from exosome (Rrp6)-depleted RNA-seq data from S2 cells (Lim et al.). Reads were adapter and quality-trimmed using cutadapt (Martin, 2011) and sickle (http://github.com/najoshi/sickle), with standard options for single-end reads. Trimmed reads from exosome knockdown samples were pooled to increase the likelihood of detecting transcripts and assembled into transcripts using Cufflinks (v2.2.1) (Trapnell et al., 2012) with non-default parameters–min-frags-per-transfrag 5 and –overlap-radius 100 for the identification of low-abundant transcripts. The transcripts were paired by overlapping their 5’ends with windows −100 to +200 around CAGE summit positions associated with each strand window of transcribed DHSs.

### Evaluation of enhancer potential by STARR-seq data

We made use of available wiggle track STARR-seq data (Zabidi et al., 2015) to evaluate the enhancer potential of transcribed DHSs. For each DHS, the summit signal within a 401 bp region centred on the DHS centre point was identified from STARR-seq data generated using *RpS12* and *even skipped* core promoters, for housekeeping (hkCP) and developmental (dCP) enhancer potential, respectively. At the summit position, we calculated the log_2_ fold change of STARR-seq signal over STARR-seq input signal. In cases in which in which analyses required distinguishing between STARR-seq positive and negative loci, STARR-seq active regions was defined as those DHSs having a log_2_ fold change of at least 1.5.

### Core promoter element scans and clustering

Core promoter element occurrences were scanned around each transcribed DHS using MEME FIMO (v4.11.2) (Grant et al., 2011). A genome sequence database of +/—50 bp around major and minor strand CAGE summit positions within each DHS was considered. Position weight matrices (PWMs) of MTE and TATA core promoter elements were retrieved from JASPAR POLII database (Mathelier et al., 2016), species *D. melanogaster.* DRE, Inr, Trl, E-box were retrieved from DMMPMM *D. melanogaster* motif collection (Kulakovskiy et al., 2009). Finally, DPE, Ohler6 (Motif6), and Ohler 1 (Motif1) motifs were collected from Ohler et al. (2002). Motif PWMs and consensus sequences were converted into the Minimal MEME Motif Format using the tools chen2meme and iupac2meme, respectively. FIMO scans were performed with a statistical threshold (p-value) equal to 1 and a maximum number of motif occurrences retained in memory at 100 × 10^6^. The FIMO output was filtered to only contain the motif hits with maximum score for each motif occurrence for each DHS strand window. Motif hits were subsequently considered significant if they passed a p-value threshold < 0.001. Core promoter element clustering was performed using the R function pheatmap (Ward.D agglomeration) on scaled and centred data for each core promoter element. Core promoter clusters were determined by the cutree R function with k=10 desired groups.

### Analyses of histone modifications and transcription factor binding at DHSs

Binarised (Zhou and Troyanskaya, 2016) modENCODE ChIP-chip data in 50 bp regions for histone modifications, histone variants and transcription factor binding was investigated around transcribed DHSs. In order to generate the heatmap against classes, the 50 bp surrounding the centre of the DHS was overlapped with the 50 bp ChIP-chip regions and the mark was recorded as present at that DHS if it overlapped with at least one element. For ChIP-chip footprint plots, the binary signal was averaged across sites in 50 bp bin intervals from the CAGE summits of the considered DHSs, up to a maximum of 5000 bp away from the summits. Cases where a given interval for a DHS overlapped another DHS were filtered from the analysis. The background level was generated based on randomising the ChIP-chip locations, 10 times for each combination of set and mark.

Chip-seq data (Herz et al. (2012)) for histone modification H3K4me3, H3K4me1 and H3K27ac were processed by the AQUAS Chip-seq pipeline (http://github.com/kundajelab/). Mapped Chip-seq signal was then quantified within each 401 bp DHS region. Chip-seq signal within DHS windows was plotted as a function of housekeeping (hkCP) or developmental (dCP) enhancer potential, using the binned 1 − 99^*th*^ percentile STARR-seq signal. Chip-seq signal within DHS windows was also plotted against the binned 1 − 99^*th*^ percentile *log*_10_ scaled CAGE TPM expression.

### Assessment of the relationship between chromatin architecture and type of regulatory element

TADs and significantly interacting regions for *D. melanogaster* Kc167 cells based on 1kb resolution HIC data (Cubeñas-Potts et al., 2017) were considered. For all HiC and TAD-associated analyses, the coordinates for +/−200 bp around the CAGE summits at DHSs were lifted over to dm6, keeping all DHSs whose width was preserved in the liftover coordinates (9,454, corresponding to 99.8% of DHSs defined in dm3). DHSs were allocated a TAD number if they overlapped the TAD by at least 200 bp. All pairs of TREs within a maximum distance of 1 Mbp between the DHS centre points were identified and annotated according to DHS class membership and whether the pair overlapped coordinates of significantly interacting regions.

For each DHS class, the proportion of elements annotated as falling inside of a TAD region, or between (not overlapping a TAD) was calculated. For each TAD, the number of DHSs of each class was aggregated. To calculate the enrichment of elements according to TAD size, TADs were split according to the total number of DHSs that were within them, grouping all TADs with more than 6 elements, and the proportion of each of the classes calculated per total size. The *log*2 scaled data containing the number of TREs per class in each TAD, for a minimum TAD size of 3 elements was further clustered using the kmeans++ algorithm, generating 7 clusters of TADs (as determined based on inspection of a scree plot for 2 to 20 possible clusters). To calculate enrichments of TAD boundary vicinities of DHS classes, a cut-off of 1kb from the nearest TAD boundary was applied to determine inclusion or exclusion of a class element from a boundary region.

Statistics of interactions and co-occurrences within TADs were analysed using generalised linear models (see Supplementary Methods).

### Statistics and visualisation

Most statistical tests and analyses were carried out in R (http://www.R-project.org). Most plots were generated using the ggplot2 R package. Intersections or distances between genomic features were calculated either using BEDTools (Quinlan and Hall, 2010) or the R package GenomicRanges (Lawrence etal., 2013).

## Acknowledgements

We thank Neus Visa for advice and sharing of reagents for the exosome knockdowns. Work in the R.A. laboratory was supported by the Danish Council for Independent Research (grant 6108-00038B) and the European Research Council (ERC) under the European Union’s Horizon 2020 research and innovation programme (grant 638173). Work in the T.H.J. laboratory was supported by the ERC (grant 339953) and the Danish National Research Council. M.L.L. was supported by the Danish Council for Independent Research (grant 1333-00059B).

## Author contributions statement

R.A. and T.H.J. conceived the project. M.L-L. and C.D.V. conducted the experiments. M.L-L. performed the experimental validations. S.R, M.D, S.B, and R.A analysed the data. S.R, M.D and R.A wrote the manuscript with input from all authors. All authors reviewed the final manuscript.

## Supplementary Materials

### Methods Supplementary Files

1. Expression and STARR-seq data associated with transcribed DHSs For each transcribed DHS, the supplementary file holds the DHS ID (hotspot coordinate), the expression (TPM) per replicate in control and exosome knockdown CAGE libraries, the exosome sensitivity on minus and plus strands, the transcriptional directionality based on control and knockdown CAGE data, the hkCP and dCP STARR-seq signal, and the DHS class.
2. Core promoter elements of transcribed DHSs For each transcribed DHS and strand, the supplementary file holds the DHS ID (hotspot coordinate) and strand, as well as the maximum motif match score and associated p-value for each considered core promoter element (DPE, DRE, E-box, INR, Ohler1, Ohler6, MTE, TATA, Trl).
3. ChIP-seq signal for transcribed DHSs For each transcribed DHS, the supplementary file holds the DHS ID (hotspot coordinate) and aggregate ChIP-seq signals for H3K27ac, H3K4me1, and H3K4me3.
4. Normalised ChIP-seq signal for transcribed DHSs For each transcribed DHS, the supplementary file holds the DHS ID (hotspot coordinate) and the counts-per-million (CPM) normalised aggregate ChIP-seq signals for H3K27ac, H3K4me1, and H3K4me3.

## Supplementary Methods

### Estimation of CAGE genomic background noise

For robust assessment of lowly expressed loci, CAGE genomic background noise levels were estimated. First, we calculated the CAGE mappability of the dm3 reference genome, by mapping each 25-sized subsequence of the reference genome back to itself, using the same mapping approach as for real CAGE data. Then, we quantified the number of CAGE 5’ ends from each out of the four control CAGE libraries mapping to genomic regions of size 200 bp, that were uniquely mappable (as determined by the mappability track) in at least 50% of its potential TSS positions (unique bps). We discarded regions that were proximal (within 500bp) of FlyBase gene TSSs, FlyBase transcript ends, or midpoints of DHS hotspot regions (as defined above), or proximal (within 100bp) of FlyBase gene exons. Based on the empirical distribution of CAGE expression noise from annotation-distal genomic regions, we extracted the 99th percentile and used the max value across control libraries as a threshold to call regions significantly expressed in subsequent analyses.

### CAGE tag clustering

First, individual genomic bps supported by at least two CAGE 5’ ends of CAGE reads in a single library that were located within 20 bp from each other on the same strand were merged into tag clusters (TCs). For each TC, the summit position (representing the within-TC bp with the highest frequency of CAGE 5’ ends) or the median of multiple summit positions was identified. Second, for each TC, the fraction between the CAGE tag frequency of each TC-covered bp to that of the summit position was calculated. All positions within a TC associated with less than 1/10 of the summit signal was discarded. This strategy causes multi-peak TCs to be split into new TCs representing each sub peak. Also, it removes tails from wide TCs with a single peak. Subsequently, TCs split on summit fraction on the same strand were merged if positioned within 20bp from each other. This resulted in a set of 670,681 TCs.

### DHSs as focus points for transcription initiation

DHSs were used as focus points for characterising patterns of transcription initiation events at TREs as described elsewhere (Andersson et al., 2014b), with minor modifications. Instead of focusing on DHS positions that maximised DNase-seq signal (DHS summits), we defined centre points of DHSs as positions optimising the coverage of proximal CAGE tags within 200 bp immediately flanking windows associated with minus and plus strand expression as illustrated in Supplementary Fig. S4. We required DHSs to be supported by significant expression above inferred noise threshold (see above) on at least one strand and in at least two out of four replicates in either control of exosome knockdown CAGE data. This resulted in 9,471 transcribed DHSs out of the original set of 11,947 DHSs. For these, exosome sensitivity and transcriptional directionality measures were calculated as defined previously (Andersson et al., 2014b). For each DHS, we defined the major and minor strand as the strand with the highest and lowest average expression level in exosome knockdown CAGE libraries, respectively.

### Annotation-unbiased clustering of DNase I hypersensitive sites

Unsupervised clustering was performed on the basis of five variables: the major and minor strand expression levels of the knockdown, the major and minor strand exosome sensitivity and the transcriptional directionality based on the knockdown samples. A total of 37,884 data vectors (4 replicates across 9,471 DHSs) were clustered. In order to determine the optimal number of clusters, we applied the kmeans++ algorithm and Mclust (Fraley and Raftery, 2002) at varying cluster sizes, with both methods concluding that either 5 or 6 clusters were optimal (based on a scree plot for cluster sizes ranging from 2 to 20 for kmeans++, and the default internal metrics from the Mclust package). We then applied the kmeans++ algorithm, which optimises the centre points of the initial centroids in the kmeans algorithm to achieve stable clusterings at 5 or 6 clusters. Correspondence analysis between pairs of replicates was used to check the overall performance of the clustering in grouping together replicates of the same DHS within a single cluster, suggesting the replicates in general to be of high quality agreement (Supplementary Fig. S5). In order to obtain a set of ‘centralised’ clusters for the 9,471 DHSs that agreed according to the clusters selected for the individual replicates, we then applied a further clustering step on the replicate cluster assignments. A distance matrix based on the hamming distance between each pair of DHSs was clustered using k-medoids, selecting for either 5 or 6 clusters. Based on biological interpretations of the resulting clusters according to their transcriptional properties, we manually choose the set based on 6 clusters, with the difference between the two clusterings surmounting to whether the *weak bidirectional unstable* and the *intermediate bidirectional unstable* DHSs should be treated as a single class or two separate classes.

### Assessment of the relationship between chromatin architecture and type of regulatory element

All pairs of transcribed DHSs within 1 Mbp were annotated according to whether they were supported by a significant interaction or not. The resulting dataset was then distance balanced, by splitting the range of distances between the pairs into 100, and for each split the number of interacting pairs was balanced against the number of non-interacting pairs (1-1 ratio, taking 10 the minimum of interaction or non-interacting). Only pairs at above the 5th split were considered, to remove the bias from difficulties of interaction calling algorithms to detect interactions at short distances. Next, pairs of transcribed DHSs were split according to pairs which overlapped dCP enhancers on both ends, pairs which overlapped hkCP enhancers on both ends or pairs which overlapped a dcP enhancer in one and a hkCP enhancer on the other element. Cases where one side overlapped both were removed from the analysis. For each class and STARR-seq enhancer class at targets, we fit a GLM to predict which of the 5 potential classes the given class interacted with. For interpretation of coefficients we generated a background class according to a random subset based on 1/5th of the dataset. We repeated the distance balancing and random background sampling 100 times, and averaged the final coefficients and p-values.

Enrichment within and between TADs proceeded similar to the GLM approach with interactions, except the response was “within TAD vs between TAD” and there was no minimum distance cut off.

**Figure S1.**
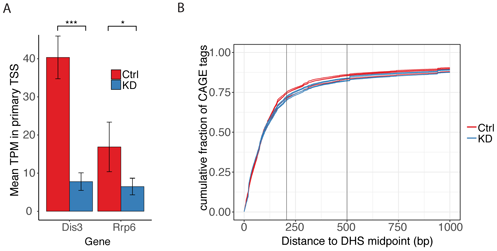
Expression of knockdown target genes and assessment of cumulative CAGE tag fractions around DHSs. **A**: Mean TPM expression of the exosome knockdown targets *Dis3* and *Rrp6* primary TSSs, measured for each control (Ctrl) and knockdown (KD) samples. Unpaired t-test determined significant reduction in knock down samples of both genes. Significance stars interpreted as: * = *P <* 0.1,*** = *P* < 0.001. **B**: Cumulative CAGE tag fraction (vertical axis) in control samples (four replicates) and knockdown samples (four replicates), as a function of the distance to the midpoints (signal summits) of DNase I hypersensitive sites (DHSs).

**Figure S2.**
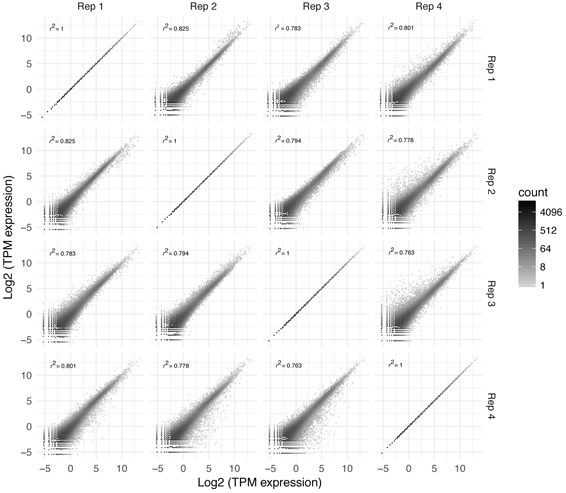
Replicate expression in control samples. Log_2_ TPM expression of TCs expressed above background noise for each pair of control replicates. Colour scale indicates number of TCs. Expression correlation between each replicate pair is indicated by r^2^ in the top left corner, based on a linear model fit.

**Figure S3.**
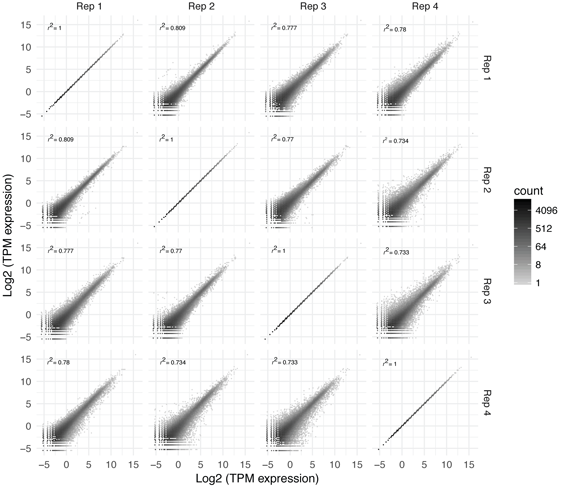
Replicate expression in knockdown samples. Log_2_ TPM expression of TCs expressed above background noise, for each pair of knockdown replicates. Colour scale indicates the number of TCs. Expression correlation between each replicate pair is indicated by r^2^ in the top left corner, based on a linear model fit.

**Figure S4.**
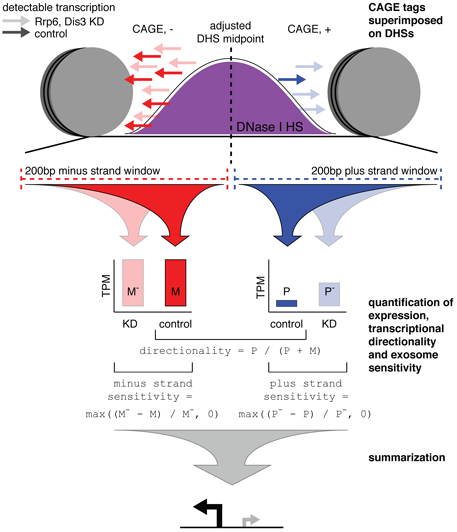
Schematic illustration of the utilisation of DNase I hypersensitive sites as focus points for transcription initiation events at regulatory elements. DHS-associated strand-specific expression levels in control and exosome depleted S2 cells were quantified by counting of CAGE tags in genomic windows of 200 bp immediately flanking the DHS centre points that optimised CAGE tag coverage (see Methods). Based on strand-specific expression levels both a directionality score, measuring the strand bias in expression level, and a strand-specific exosome sensitivity score, measuring the relative amount of degraded RNAs by the exosome, were calculated. These measures were used for clustering the DHSs to infer classes of DHSs with similar transcriptional properties (Fig. 2A).

**Figure S5.**
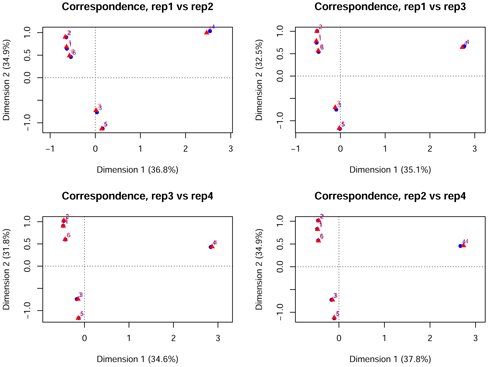
Correspondence analysis of CAGE replicate agreement in DHS clustering.

**Figure S6.**
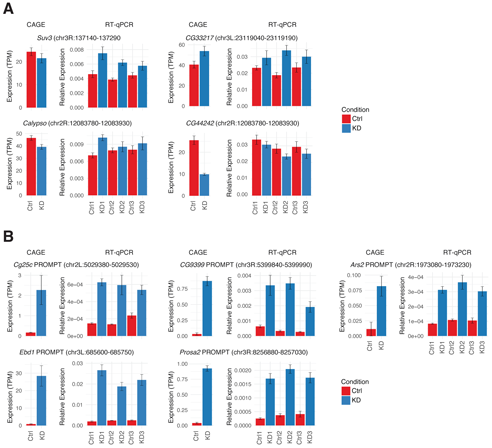
CAGE expression and RT-qPCR validation of mRNA and PROMPT tagets. **A:** CAGE expression and RT-qPCR validation of four randomly selected *unidirectional stable* targets, listed with FlyBase gene ID and DHS coordinates. For each target, left panel shows mean CAGE expression (TPM) and standard deviations of DHS major strand (gene sense direction), within all four replicates per CAGE condition; control (Ctrl) and knock down (KD). Right panel shows RT-qPCR mean relative expression with standard deviations of three technical replicates (normalised to *Act5C*), for three biological replicates per condition; control (Ctrl) and knock down (KD). **B**: CAGE expression and RT-qPCR validation of four randomly selected PROMPTs of the *unidirectional stable w*/ *PROMPT* DHS cluster, together with the *Cg25C* PROMPT shown in Fig. 1A. FlyBase gene ID and DHS coordinates are listed on top of each PROMPT plot section. For each target, left panel shows mean CAGE expression (TPM) and standard deviations of DHS minor strand (antisense of gene direction), within all four replicates per CAGE condition; control (Ctrl) and knock down (KD). Right panel shows RT-qPCR mean relative expression of PROMPT targets (upstream of and antisense to annotated FlyBase gene TSSs), with standard deviations of three technical replicates (normalised to *Act5C),* for three biological replicates per condition; control (Ctrl) and knock down (KD).

**Figure S7.**
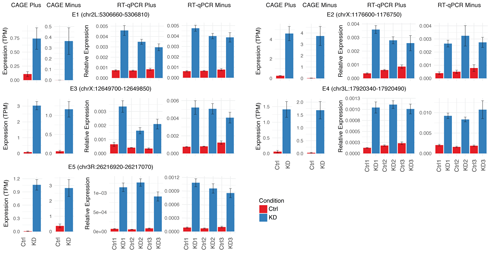
CAGE expression and RT-qPCR validation of bidirectional unstable transcription. CAGE expression and RT-qPCR validation of four randomly selected enhancer RNA tagets from the *bidirectional weak unstable* DHS cluster, together with the intragenic dCP STARR-seq enhancer shown in Fig. 1B. DHS coordinates are listed above each plot section. For each target, left panel shows DHS plus and minus strand mean CAGE expression (TPM) and standard deviations within all four replicates per CAGE condition; control (Ctrl) and knock down (KD). Right panel shows RT-qPCR mean relative expression with standard deviation of three technical replicates (normalised to *Act5C)* for three biological replicates per condition; control (Ctrl) and knock down (KD). RT-qPCR was performed on both plus and minus TC peak

**Figure S8.**
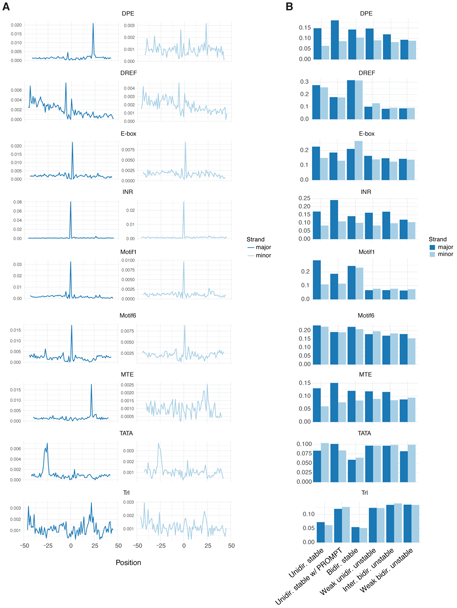
Fraction of core promoter elements on minor and major strands of transcribed DHSs. **A**: Fraction of transcribed DHSs (vertical axis) with an identified core promoter element at a given position relative to the major (left panels) and minor (right panels) strand CAGE summits. **B**: Fraction of DHSs with a core promoter element in each DHS class broken up by major and minor strand.

**Figure S9.**
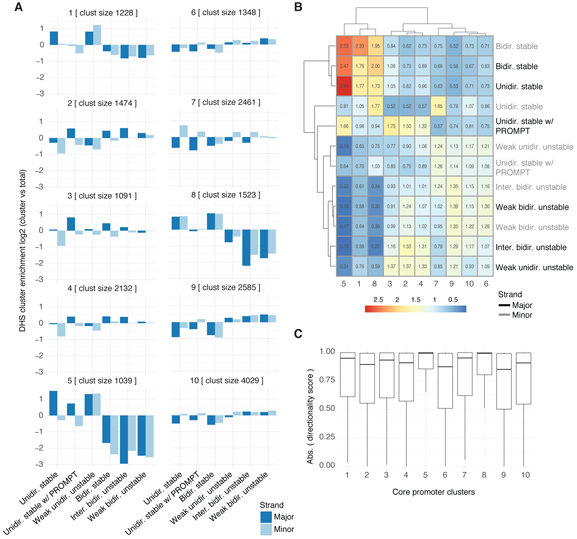
DHS enrichment for identified core promoter element clusters. **A**: DHS class enrichments, calculated as the fraction of DHSs in each DHS class associated with each core promoter element cluster versus the fraction of total transcribed DHSs, displayed in log_2_ scale enrichment, broken up by major and minor strand. **B**: DHS class enrichment, hierarchically clustered with complete linkage, for major and minor strands separately, within each identified core promoter element clusters. **C**: Absolute directionality score within each identified core promoter cluster.

**Figure S10.**
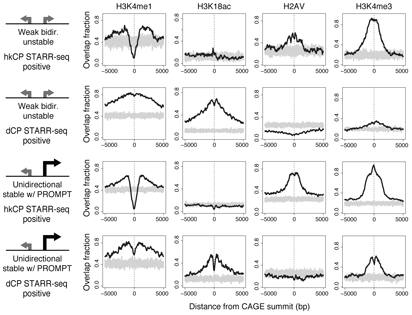
Binding enrichments at DHS classes with respect to enhancer potential. Detailed binding enrichments for H3K4me1, H3K18ac, H2Av, and H3K4me3 at *weak bidirectional unstable* and *unidirectional stable w/ PROMPT* DHSs, broken up according to STARR-seq enhancer potential (overlapping either a hkCP or dCP enhancer), based on binding proportions within 5,000 bp from the CAGE summit. Grey represents background distribution based on randomised locations, generated 10 times per plot.

**Figure S11.**
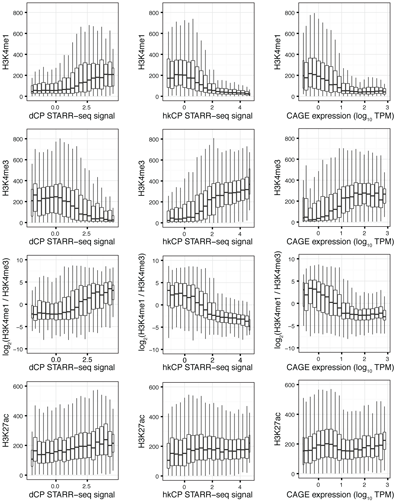
Histone modifications versus STARR-seq signal and expression strength. Normalised ChIP-seq data of H3K4me1, H3K4me3, log_2_ H3K4me1 over H3K4me3 ratio, and H3K27ac (vertical axis) versus binned dCP (left) and hkCP (middle) STARR-seq signal, and binned TPM scaled log_10_ CAGE expression (right) (horisontal axes).

**Figure S12.**
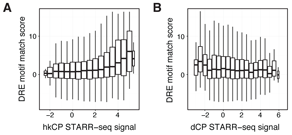
DRE motif score versus STARR-seq signal. FIMO DRE motif match score (see Methods) versus STARR-seq log_2_ signal fold change versus input for hkCP (**A**) and dCP (**B**) enhancer potential. STARR-seq signal is binned in 0.5 sized ranges.

**Figure S13.**
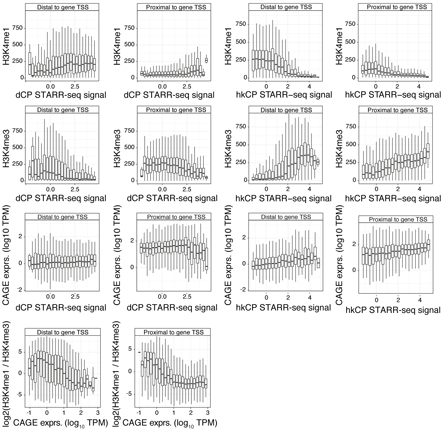
Histone modifications versus STARR-seq signal and expression strength of gene proximal and distal DHSs. Normalised ChIP-seq data of H3K4me1, H3K4me3, log_2_ H3K4me1 over H3K4me3 ratio, and TPM scaled expression (vertical axis) versus binned dCP (left) and hkCP (right) STARR-seq signal, and binned TPM scaled log_10_ CAGE expression (bottom) (horisontal axes).

**Figure S14.**
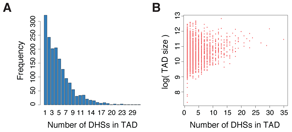
DHSs linked to TAD information. **A**: The frequency of transcribed DHSs within Kc167 TADs. **B**: Number of DHSs within a TAD versus the size of the TAD in which they were contained.

**Figure S15.**
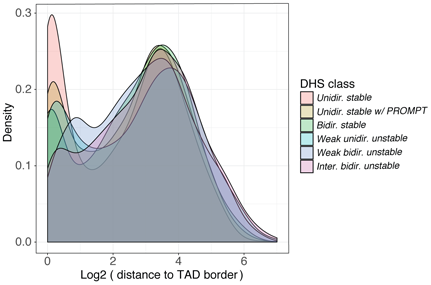
DHS distance to TAD borders. Smoothed density estimates of log_2_ scaled kb distances per DHS class, measured between DHS windows within Kc167 TADs and TAD borders.

**Figure S16.**
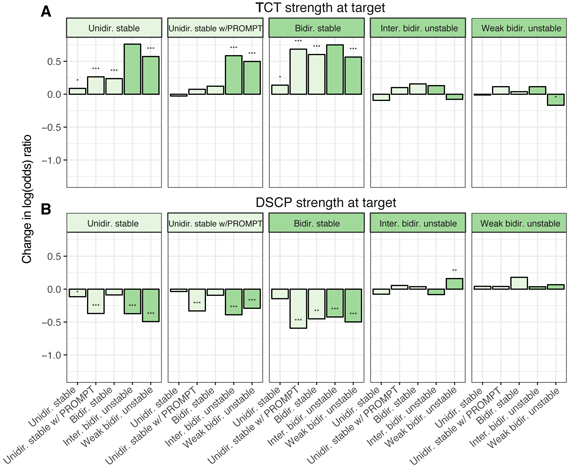
STARR-seq vs DHS class interaction preferences. (**A-B**):Class chromatin-interaction preferences based on modelled interactions with STARR-seq strength overlap (see Methods). Significance stars interpreted as: * = *P <* 0.1, ** = *P* < 0.01 or *** = *P* < 0.001. A single model was fitted per bait class, with all covariates together which were scaled prior to model fitting. **A**: Modelled interaction between target class and hkCP enhancer strength (log_2_(signal over input)) at the target. **B**: Modelled interaction between target class and dCP enhancer strength (log_2_(signal over input)) at the target.

**Table S1.** Cluster agreement statistics from k-medoids clustering of replicate cluster assignments based on the hamming distance between each pair of DHSs (max_diss: max dissimilarity, av_diss: average dissimilarity).

| DHS class | size | max_diss | av_diss | diameter | separation |
| --- | --- | --- | --- | --- | --- |
| Weak bidir. unstable | 1723 | 3 | 0.85 | 4 | 1 |
| Unidirectional stable w/ PROMPT | 1633 | 3 | 1.00 | 4 | 1 |
| Weak unidir. unstable | 1232 | 2 | 0.82 | 4 | 1 |
| Bidirectional stable | 912 | 2 | 0.14 | 4 | 1 |
| Intermediate bidir. unstable | 1556 | 3 | 0.88 | 4 | 1 |
| Unidirectional stable | 2415 | 2 | 0.40 | 4 | 1 |

**Table S2.** RT-qPCR relative mean expression. Mean relative expression of three technical replicates (normalised to *Act5C*) for all validated target sites in three biological replicates, listed by DHS coordinates and FlyBase gene name, where P indicates PROMPT target upstream and antisense of gene, or enhancer ID (E) including TC strand information.

| DHS ID | Name | Ctrl 1 | KD 1 | Ctrl 2 | KD 2 | Ctrl 3 | KD 3 |
| --- | --- | --- | --- | --- | --- | --- | --- |
| 3R:137140-137290 | *Suv3* | 4.605e-03 | 7.49e-03 | 3.854e-03 | 6.201e-03 | 4.445e-03 | 5.764e-03 |
| 3L:23119040-23119190 | *CG33217* | 0.023219 | 0.029347 | 0.018738 | 0.033844 | 0.02343 | 0.030002 |
| 2R:12083780-12083930 | *Calypso* | 7.038e-03 | 0.010163 | 7.921e-03 | 8.561e-03 | 8.036e-03 | 9.162e-03 |
| 2R:12083780-12083930 | *CG44242* | 0.033783 | 0.03073 | 0.028301 | 0.023585 | 0.02938 | 0.02526 |
| 2L:5029380-5029530 | *Cg25cp* | 1.46e-04 | 6.25e-04 | 1.35e-04 | 5.93e-04 | 2.38e-04 | 5.36e-04 |
| 3R:5399840-5399990 | *CG9399p* | 6.36e-04 | 3.35e-03 | 3.29e-04 | 3.479e-03 | 2.72e-04 | 1.919e-03 |
| 2R:1973080-1973230 | *Ars2p* | 8.3e-05 | 3.13e-04 | 1.08e-04 | 3.64e-04 | 1.04e-04 | 3.03e-04 |
| 3L:685600-685750 | *Ebd1p* | 1.913e-03 | 0.026606 | 2.35e-03 | 0.018791 | 2.468e-03 | 0.021944 |
| 3R:8256880-8257030 | *Prosa2p* | 2.49e-04 | 1.697e-03 | 3.71e-04 | 2.045e-03 | 4.12e-04 | 1.73e-03 |
| 2L:5306660-5306810 | *E1plus* | 7.4e-04 | 4.581e-03 | 7.12e-04 | 3.491e-03 | 8.37e-04 | 2.96e-03 |
| 2L:5306660-5306810 | *E1minus* | 6.28e-04 | 4.747e-03 | 6.57e-04 | 3.997e-03 | 7.78e-04 | 3.875e-03 |
| X:1176600-1176750 | *E2plus* | 3.64e-04 | 3.588e-03 | 6.08e-04 | 2.796e-03 | 8.61e-04 | 2.587e-03 |
| X:1176600-1176750 | *E2minus* | 3.9e-04 | 2.648e-03 | 4.98e-04 | 3.24e-03 | 7.81e-04 | 2.742e-03 |
| X:12649700-12649850 | *E3plus* | 6.52e-04 | 3.354e-03 | 4.15e-04 | 1.626e-03 | 3.49e-04 | 2.101e-03 |
| X:12649700-12649850 | *E3minus* | 7.59e-04 | 5.212e-03 | 8.05e-04 | 5.071e-03 | 1.233e-03 | 4.064e-03 |
| 3L:17920340-17920490 | *E4plus* | 1.28e-04 | 1.041e-03 | 1.79e-04 | 1.107e-03 | 2.32e-04 | 1.008e-03 |
| 3L:17920340-17920490 | *E4minus* | 2.052e-03 | 9.212e-03 | 1.599e-03 | 8.322e-03 | 1.889e-03 | 0.010724 |
| 3R:26216920-26217070 | *E5plus* | 6.8e-05 | 1.147e-03 | 5.3e-05 | 1.27e-03 | 8e-05 | 9.08e-04 |
| 3R:26216920-26217070 | *E5minus* | 8.8e-05 | 1.082e-03 | 7e-05 | 9.55e-04 | 9.9e-05 | 8.52e-04 |

**Table S3.**
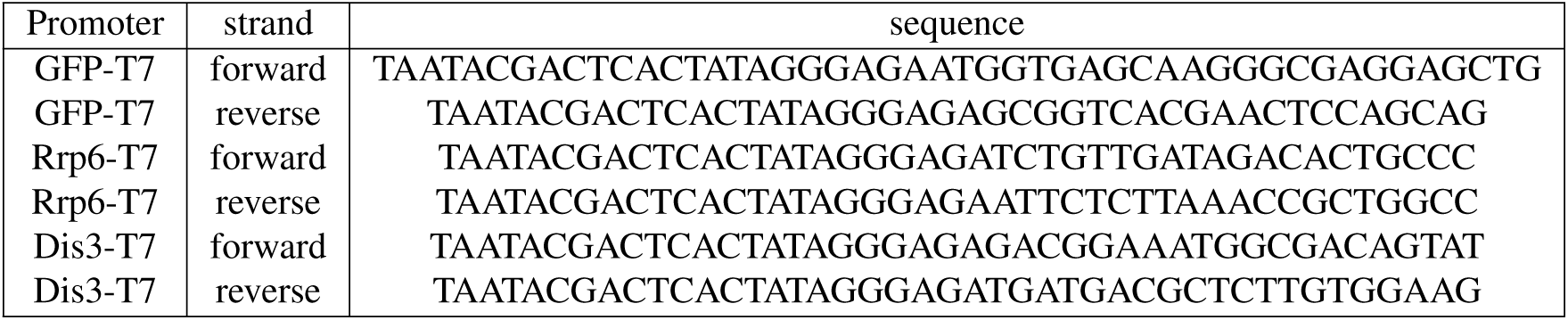
Oligos for PCR templates for making dsRNA.

**Table S4.**
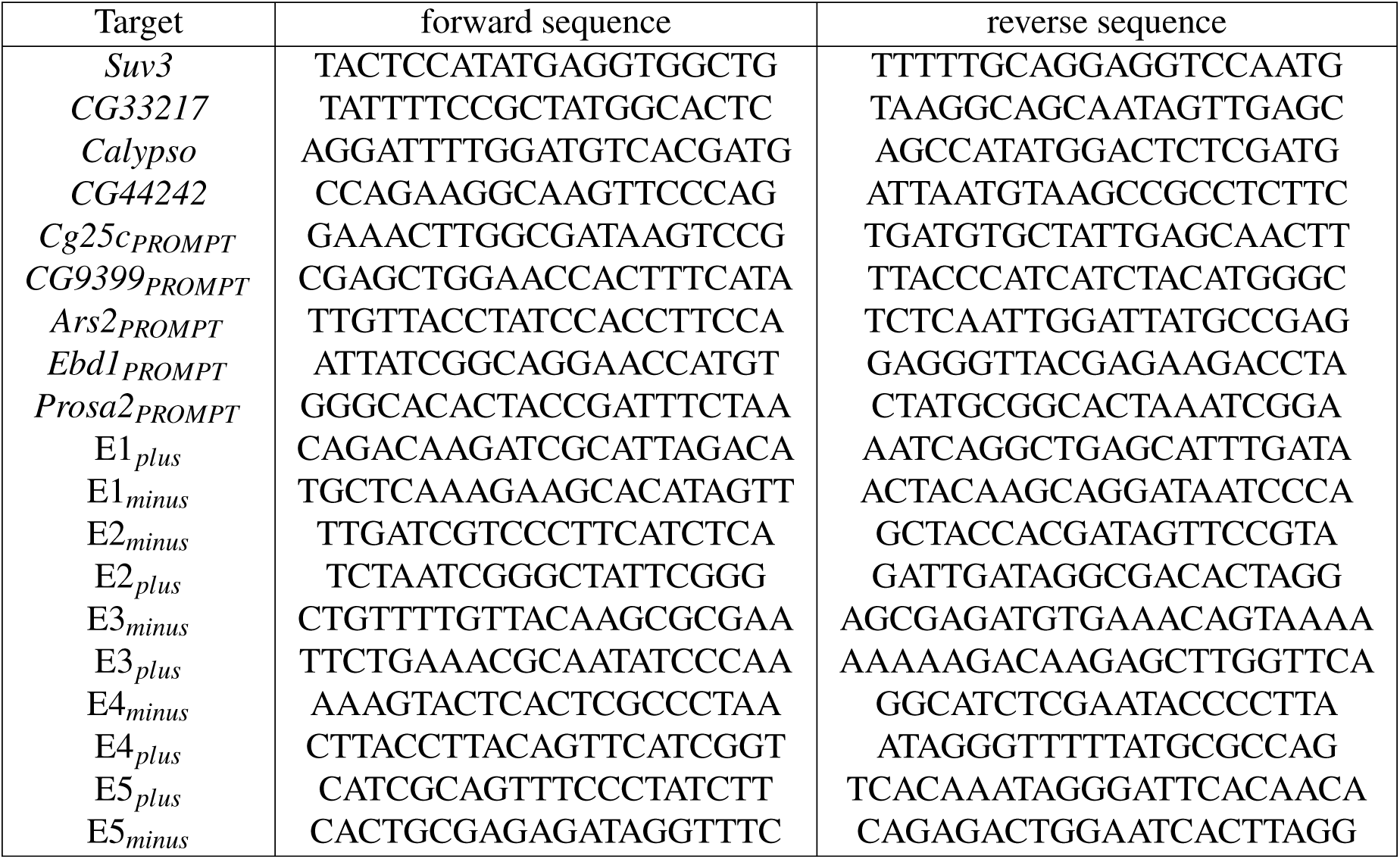
Oligos for qPCR.

